# TLR4 signaling in neurons enhances calcium-permeable AMPAR currents and drives post-traumatic epileptogenesis

**DOI:** 10.1101/649780

**Authors:** Akshata A. Korgaonkar, Ying Li, Dipika Sekhar, Deepak Subramanian, Jenieve Guevarra, Bogumila Swietek, Alexandra Pallottie, Sukwinder Singh, Kruthi Kella, Stella Elkabes, Vijayalakshmi Santhakumar

**Author notes:** Correspondence: Akshata Korgaonkar, PhD, Department of Neurology, Washington University School of Medicine, 660 South Euclid Ave, Campus box 8111, St Louis, MO 63110, Phone (Off).

## Abstract

Traumatic brain injury is a major risk factor for acquired epilepsies and understanding the mechanisms underlying the early pathophysiology could yield viable therapeutic targets. Growing evidence indicates a role for inflammatory signaling in modifying neuronal excitability and promoting epileptogenesis. Here, we identify that signaling through an innate immune receptor, toll-like receptor 4 (TLR4), in neurons, augments calcium-permeable AMPA receptor (CP-AMPAR) currents in the hippocampal dentate gyrus after brain injury. Blocking TLR4 signaling *in vivo* shortly after brain injury reduced dentate network excitability and seizure susceptibility. When blocking of TLR4 signaling after injury was delayed, however, this treatment failed to reduce post-injury seizure susceptibility. Further, TLR4 signal blocking was less efficacious in limiting seizure susceptibility when AMPAR currents, downstream targets of TLR4 signaling, were transiently enhanced. Paradoxically, blocking TLR4 signaling augmented both network excitability and seizure susceptibility in uninjured controls. Despite the differential effect on seizure susceptibility, TLR4 antagonism suppressed cellular inflammatory responses after injury without impacting sham controls. These findings demonstrate that independently of glia, the immune receptor TLR4 directly regulates post-traumatic neuronal excitability. Moreover, the TLR4-dependent early increase in dentate excitability is causally associated with epileptogenesis. Identification and selective targeting of the mechanisms underlying the aberrant TLR4-mediated increase in CP-AMPAR signaling after injury may prevent epileptogenesis after traumatic brain injury.

**Graphical Abstract:** *Summary of interactions between TLR4 signaling and brain injury on network excitability and epileptogenesis:* Graphic illustration of the effect of injury and early TLR4 antagonist treatment on early network excitability and the long-term network state. The schematic neurons include TLR4 and AMPAR subunit expression profiles in the acute phase of sham or brain injury. The corresponding early effects on network excitability are depicted by schematic population response traces (inset on upper left). Note the increase in excitability of the uninjured neuron after TLR4 antagonism without changes in AMPAR expression. Note also the increase in TLR4, calcium permeable AMPARs and population excitability after injury and its reduction by TLR4 antagonist treatment. Ampakine enhancement of excitability during TLR4 antagonism is illustrated. The early phase responses and manipulations (including injury, treatments, and molecular responses) are superimposed on a two-tone color-coded network state topology where green indicates low-normal network excitability, ensuring network stability and low risk for epilepsy (Inset on upper right). Note the correspondence between early excitability state (population response profile) and long-term seizure susceptibility and the effects of pharmacological manipulations.

## Introduction

Acquired epilepsies that develop after brain insults such as trauma are particularly refractory to treatments yet are potentially preventable if the underlying mechanisms are identified and appropriately targeted (1–3). A wealth of preclinical and clinical studies predict that neuronal excitability and plasticity act alongside inflammatory processes and contribute to epileptogenesis (3–9). However, the interaction between neurophysiological and inflammatory responses to injury, and the underlying mechanisms are not fully understood (10).

Activation of the innate immune receptor toll-like receptor 4 (TLR4) by its ligand HMGB1, which is released during neuronal damage, is thought to play a critical role in the inflammatory responses to seizures and brain injury (6, 11). TLR4 is expressed in both neurons and glia (11, 12). Glial TLR4 signaling has been proposed to underlie increases in neuronal excitability and excitotoxicity by activating glial cytokines, which enhance neuronal NMDA receptor (NMDAR) currents (11, 13, 14). In contrast, we recently identified TLR4 expression and enhancement in hippocampal dentate neurons after experimental brain injury, but not in astrocytes or microglia (12). Curiously, acute ex vivo antagonism of TLR4 signaling modulated AMPA receptor (AMPAR), but not NMDAR currents, in dentate neurons in slices one week after brain injury. These data raise the possibility that TLR4 signaling after brain injury does not engage the classical glial pathways (15), but instead acts through neuronal effectors which are yet to be identified (16, 17). The ability of TLR4 antagonists to reduce early network excitability in slices (12) raises the untested, clinically relevant, possibility that TLR4 signaling contributes to and could be harnessed to limit epileptogenesis after brain injury. Moreover, it is imperative to identify whether the paradoxical increase in network excitability in slices from uninjured rats treated with TLR4 antagonists (12) has both immediate and lasting consequences on dentate networks in vivo in order to evaluate the feasibility of targeting TLR4 for therapeutics. The divergent effect of TLR4 on network excitability underscores the need to identify the effectors and mechanisms underlying these disparate responses. Simultaneously, differential TLR4-mediated effects present an unparalleled opportunity to address the long-standing question concerning the role for altered network excitability in post-traumatic epileptogenesis (4, 5) and to discriminate between the roles for neuronal and classical inflammatory effectors in injury-induced epileptogenesis.

Here we use a rodent fluid percussion injury (FPI) model of concussive brain injury and post-traumatic epileptogenesis to identify the cellular, synaptic and receptor mechanisms underlying TLR4 enhancement of hippocampal dentate excitability within a week after brain injury. Focusing on cellular and molecular processes in the injured brain, we demonstrate that TLR4 signaling in neurons rapidly enhances calcium-permeable AMPAR (CP-AMPAR) currents. Our data show that early changes in hippocampal dentate excitability predict risk for epileptogenesis, regardless of the presence or absence of cellular inflammation. We identify that transient and early inhibition of the TLR4-AMPAR signaling in vivo after brain injury has the potential to prevent post-traumatic epileptogenesis.

## Materials and Methods

### Animals

All procedures were approved by the Institutional Animal Care and Use Committee of the Rutgers New Jersey Medical School, Newark, New Jersey.

### Fluid percussion injury

Juvenile male Wistar rats (25-27 days old) were subject to the moderate (2.0-2.2 atm) lateral fluid percussion injury or sham-injury using standard methods (12, 18, 19). Only rats with >10 sec acute apnea and startle/seizures acutely after injury were included in the injury group and only the ipsilateral hemisphere was examined.

### Drug administration and electrode implantation

One day (20-24 hrs.) after injury, a randomly assigned cohort of FPI and sham rats received either vehicle (saline) or a synthetic TLR4 antagonist, CLI-095 (0.5mg/kg, s.c.) for 3 days. A second group underwent stereotaxic injection (Hamilton syringe-26 G) of 5µl of saline or LPS-RS Ultrapure (LPS-RS*U*, 2mg/ml) to the ipsilateral hippocampus. Injections were delivered through the implanted syringe hub, which was used to deliver FPI, one day after injury at a rate of 1µl/5 mins. Some animals received an Ampakine (CX546, 300µM) along with LPS-RS*U*. A third group of animals underwent stereotaxic placement of a cannula electrode in the ipsilateral hippocampus (18). Briefly, after injury, sham and FPI rats underwent stereotaxic placement of a cannula electrode through the implanted syringe hub. Drugs were then injected through the cannula directly to the hippocampus. Some animals underwent stereotaxic implantation of an ipsilateral hippocampal depth or surface electrode 25 days or 85 days after FPI or sham injury. Drug dosing was based on published studies *in vivo* (20–23).

### Electrophysiology

Six to eight days after FPI or sham-injury, rats were anesthetized with isoflurane and decapitated. Horizontal brain slices (400 µm for field recordings and 350 µm for whole cell recordings) were prepared as described previously (12, 24). Field recording were conducted in an interface field recording chamber (BSC2, Automate Scientific, Berkeley, CA), perfused, and recorded in ACSF at 32–33°C making note of the electrode positions. Field recordings of dentate population responses were obtained using patch pipettes containing ACSF (12, 25). Slices were then incubated in CLI-095 or ACSF for 45 min and transferred back to the recording chamber for recordings. For whole cell patch clamp recordings of afferent evoked AMPARs, currents were recorded and analyzed as detailed previously (12). CP-AMPAR current was recorded at −60 mV and +40 mV holding potential in the presence of SR95531 (10µM) and D-APV (50 µM). ClampFit 10 and SigmaPlot 12.3 were used to analyze the data.

### Cell surface protein extraction and Western blotting

The cell surface proteins of hippocampus were isolated using Pierce cell surface protein isolation kit (Pierce, Thermo Scientific, MA). Biotinylated proteins bound to NeutrAvidin beads (bound fraction) were isolated from the nonbiotinylated (unbound) fraction by centrifugation (3000 RPM, 1 min). The bound proteins were then released by incubating with Laemmli sample buffer containing 50mM of dithiothereitol. Cell surface proteins were isolated from experimental groups and the extracted proteins as well as total proteins were used for western blotting analysis. Western blots were performed as described previously (12). Membranes were then washed and incubated overnight at 4°C with the corresponding primary and secondary antibodies. Chemiluminescent detection was performed using FluoroChem 8800, and images were quantified using Image-J (NIH). Surface protein expression was normalized to corresponding total protein levels and total protein levels were normalized and normalized to β-actin density. While surface proteins did include a faint band for β-actin levels in the total protein and likely reflect proteins associated with intracellular membranes. All antibodies used in the study are in Supplementary Table 3.

### Neuronal Cultures

Primary hippocampal neurons were obtained from the hippocampi of Embryonic day 17-18 mice (*WT* or *TLR4^-/-^*) embryos using standard methods (26, 27). Hippocampal tissue was centrifuged at 1000 rpm for 1 min with Hanks balanced salt solution (HBSS), digested DNAse (200 Units/mg) in 25% trypsin solution washed with and transferred to Neurobasal media-A (NB-A) and mechanically dissociated. The cells were pelleted by centrifugation in NB-A and 0.5-1.0 x 106 cells per well were plated on Poly-D-Lysine (PDL) coated 15mm coverslips in NB-A medium containing B27 and L-glutamine and cultured at 37°C in 5% C02/95% air. The medium was changed every 3-4 days. Ara-C was added to media on day 3 for complete removal of glia. Cultured coverslips were stained for MAP2, GFAP and IBA-1 to identify cell types. 11 to 13 DIV cultures were used for patch clamp recordings described above.

### Histology

For immunostaining, rats were perfused with 4% paraformaldehyde 24hr after FPI or sham-injury and hippocampal sections (50 µm) were obtained. Immunostaining was performed as detailed previously using antibodies listed in supplementary table 3. Nissl staining was performed on coronal sections (50 µm) from experimental rats perfused with 4% paraformaldehyde 1 week after injury. Sections were mounted on gelatinized slides, air dried, stained in Cresyl violet and differentiated in alcohol and xylene prior to mounting. Cell counts were performed using the optical fractionator of Stereo Investigator V.10.02 (MBF Bioscience, Williston, VT) on an Olympus BX51 microscope with a 40X objective. In each section, the hilus was outlined by a contour traced using a 10X objective. Sampling parameters were set at 40X: counting frame = 50 µm by 50 µm, dissector height = 30 µm, and top guard zone = 5 µm. Approximately 25 sites per contour were selected using randomized systematic sampling protocols in Stereo Investigator(28).

### Video-EEG monitoring

One and three months after injury, rats were implanted with a hippocampal depth electrode as described above and seizure susceptibility to Kainic acid (KA, 5 mg/kg i.p.) was examined using a video EEG recording system (18, 29). Three to four months after brain injury a group of experimental rats were implanted with subdural screw electrodes and monitored for development of spontaneous seizures using video EEG recording system as described previously (24). Animals were monitored for 8 continuous hours overnight for 3-5 days (minimum of 5 days if seizures were not detected). Following EEG monitoring, rats were transcardially perfused with saline followed by 4% paraformaldehyde under surgical anesthesia.

### EEG data analysis

Analysis was performed by investigators blinded to treatment groups. Electrographic seizures were defined as rhythmic activity exceeding a threshold of mean baseline amplitude + 2.5 S.D. for >5 seconds. Simultaneous video recordings were used to exclude movement artifacts. For kainic acid evoked seizures, latency of first onset electrographic and behavioral seizure, and seizure severity were analyzed for 3 hours post injection. Seizure severity was assessed off-line by ranking the behavior according to a modified Racine’s scale (30): 0 = no behavioral change (subclinical), 1 = facial movements (twitching of vibrissae, sniffing, eye blinking or jaw automatisms), 2 = head nodding, stiff tail, 3 = forelimb clonus, chewing, 4 = rearing with myoclonus and tonic immobility and 5= Rearing and falling with myoclonus (tonic-clonic seizures). For spontaneous seizures, only electrical seizures associated with stage 3 or higher behavioral seizures and lasting at least 30 seconds were included in the analysis.

### Flow Cytometry

Three days after injury, animals were anesthetized, perfused with cold saline and the hippocampus from the injured hemisphere and spleen (positive staining control) extracted, dissociated and processed as detailed previously (31). Following LIVE/DEAD staining, cells were resuspended in FACS buffer and probed for CD45, CD3, CD4, GR-1, and OX42. Appropriate isotype controls were included to reduce non-specific signal (Supplementary Table 3). Samples (1,000,000 events/sample) were acquired using BD LSRII flow cytometer and data analyzed using FlowJo (V 10.0.8).

### Statistical analysis

Statistical analyses were performed using SigmaPlot 12.3. Analysis was performed once, midway during experiments to determine sample size. Data are shown as mean ± s.e.m. in Supplementary Table 1. Statistical results are presented in Supplementary Table 2.

## Results

### TLR4 signaling enhances CP-AMPAR current in dentate granule cells after brain injury

Immunostaining for TLR4 in the hippocampal dentate gyrus neurons was confirmed by co-localization of TLR4 with the neuronal marker NeuN (Fig. 1a) (12). Consistent with the effects of LPS-RS*U* (12), CLI-095 (or TAK-242/Resatorvid, 10ng/ml), an antagonist at the intracellular domain of TLR4 (32), reversed increases in afferent-evoked dentate population spike amplitude and field EPSP slope in slices one week after brain injury (Fig. 1b-e, Supplementary tables). However, CLI-095 increased excitability in slices from sham operated controls (Fig. 1c-e) demonstrating an opposite effect in the uninjured brain. Previously, we identified a selective increase in granule cell AMPAR currents one week after brain injury, with no changes in NMDAR currents (12, 33). TLR4 antagonists abolished post-injury increases in AMPAR current amplitude without altering AMPAR or NMDAR currents in controls (12). Granule cells express heteromeric AMPARs containing both GluA1 and GluA2 subunits, with the presence of GluA2 rendering the AMPAR currents calcium impermeable and non-rectifying (34, 35). In controls, pharmacologically isolated, perforant-path evoked granule cell AMPAR currents recorded from holding potentials of +40 and −60 mV were symmetrical (non-rectifying) indicating a calcium-impermeable phenotype (Fig. 1f-g). The selective TLR4 antagonist LPS-RS*U* (1µg/ml) failed to alter AMPAR current rectification in controls. In contrast, granule cell AMPAR currents were inwardly-rectifying one week after brain injury (Fig. 1f-g), indicating an increase in CP-AMPAR currents. Incubation in LPS-RS*U* reduced rectification (Fig. 1f-g), demonstrating that TLR4 signaling underlies increases in CP-AMPAR currents after injury. The molecular underpinnings of the post-injury increase in CP-AMPAR currents were examined in hippocampal slices from control and injured rats treated with saline or LPS-RS*U* (1µg/ml for 1hr). Western blots of surface biotinylated AMPAR subunits identified an increase in surface expression of GluA1 subunit after brain injury, which was reversed by a brief (1hr) incubation in LPS-RS*U* (Fig. 1h-i). LPS-RS*U* failed to alter GluA1 surface expression in controls, and GluA2 surface expression was unchanged after injury or with LPS-RS*U* treatment (Fig. 1i-j). Unlike surface GluA1 expression, total GluA1 and GluA2 expression were unchanged after FPI and with drug treatments (Fig 1i, k-l). These data demonstrate that TLR4 signaling selectively and reversibly augments membrane surface expression of the GluA1 subunit early after brain injury. The increase in GluA1 in the absence of corresponding increase in GluA2 would increase the faction of GluA2 lacking AMPARs and contribute to the post-injury increase in CP-AMPAR currents.

**Fig. 1.**
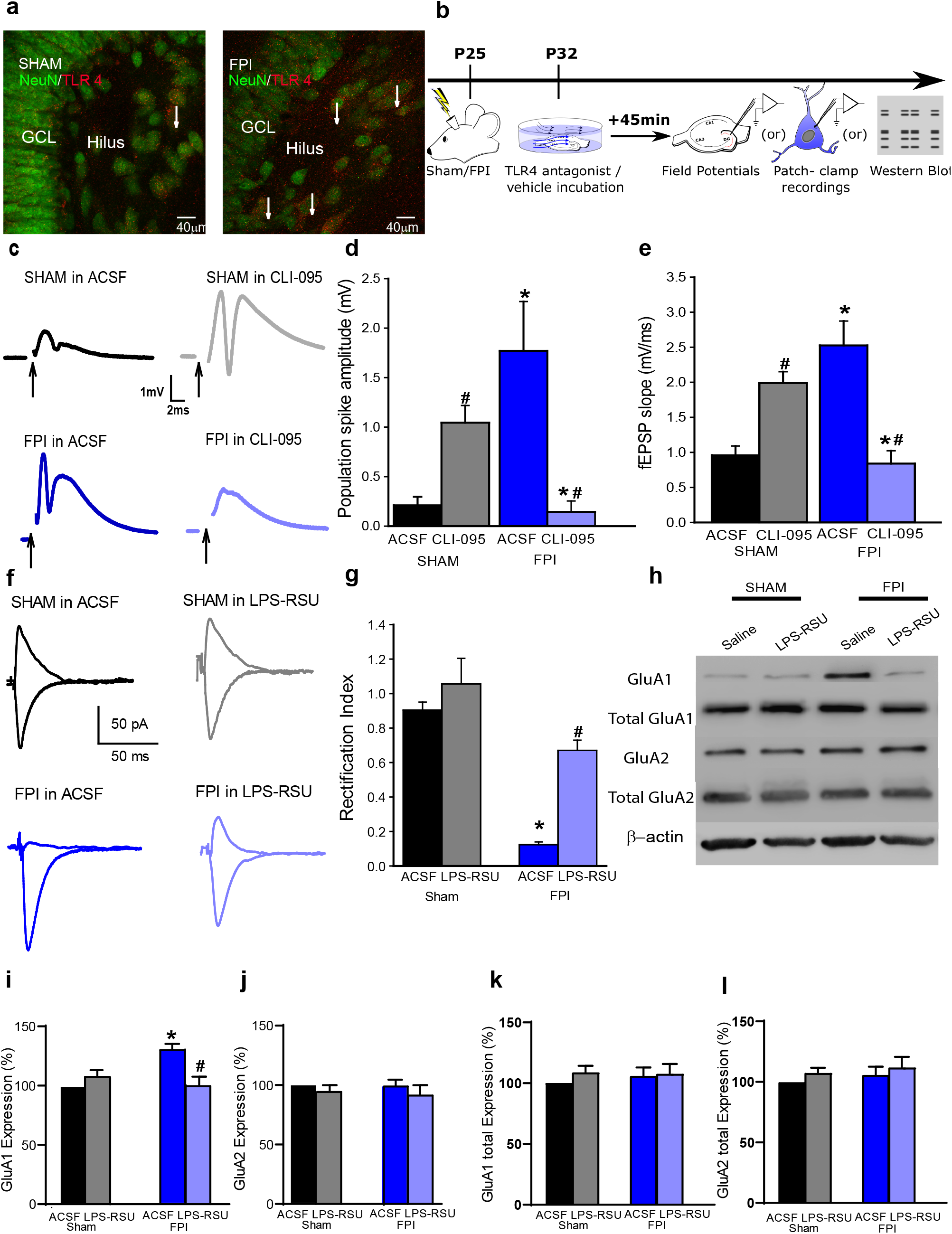
TLR4 signaling augments CP-AMPAR currents and GluA1 surface expression after brain injury. a. Representative images of dentate granule and hilar cells in rats sacrificed 24 hours after sham or FPI illustrate co-localization of TLR4 with NeuN (arrows). b. Schematic of experimental design for panels c-j. Following sham or FPI at postnatal day (P) 25, hippocampal tissue was isolated at P32 for experiments. c. Granule cell population responses to perforant path stimulation (4mA) in a slice from a sham (upper panels) and FPI rats (lower panels) incubated in ACSF (left) and CLI-095 (10 ng/ml for 1 h, right). Arrows indicate truncated stimulus artifact. d-e. Summary histogram of population spike amplitude (d) and field EPSP slope (e) in slices from rats 1-week post-injury. (* indicates p<0.05 compared to without drug; # indicates p<0.05 compared to sham by TW-RM-ANOVA followed by post-hoc Tukey’s test). f. Perforant path-evoked granule cell AMPAR current traces recorded at +40 and −60 mV, in the presence of GABA_A_R and NMDAR blockers, show lack of rectification in sham (above left) and in LPS-RS*U* (above right). One week after FPI, AMPAR currents show rectification (below left) which was reversed in LPS-RS*U* (below right). g. Summary of rectification index measured as the ratio of peak AMPAR current at +40 mV and −60 mV. (* indicates p<0.05 by TW-ANOVA). h. Western blots for GluA1 and GluA2 expression in surface fractions extracted following biotinylation assay on hippocampal sections from sham and FPI rats incubated in ACSF or LPS-RS*U*. i-l. Summary plots quantify the relative expression of surface biotinylated GluA1 (i) and GluA2 (j) expression normalized to total GluA1 and GluA2 respectively and total GluA1 (k) β-actin and presented as % of control. *p< 0.05 from sham and ^#^ indicates p<0.05 compared to saline within injury type by TW ANOVA followed by pairwise comparison.

### Neuronal signaling underlies TLR4-modulation of AMPAR currents

While dentate neurons express TLR4 (12), direct regulation of CNS neuronal excitability by TLR4 remains to be established. To isolate the effects of neuronal TLR4, neuronal cultures at day *in vitro* (DIV) 12 were established from embryonic mouse hippocampi and verified for neuronal purity by positive immunolabeling of the neuronally enriched microtubule associate protein 2 (MAP2), and lack of immunolabeling for GFAP and Iba1 (Fig. 2a-b), markers for astrocytes and microglia, respectively. In neuronal cultures from *wild-type* (*WT*) mice, neuropil stimulation evoked synaptic AMPAR currents with inward rectification, indicating calcium-permeable currents (Fig. 2c-d), similar to currents seen in granule cells after brain injury. This is consistent with reports of increased CP-AMPAR currents during development *in vitro* (36, 37). However, AMPAR currents in neuronal cultures from mice lacking TLR4 (*TLR4^-/-^*) were non-rectifying (Fig. 2c-d) demonstrating that neuronal TLR4 is necessary for expression of the inwardly rectifying CP-AMPAR currents. In neurons cultured from *WT* mice, the TLR4 agonist HMGB1 (10ng/ml) failed to increase AMPAR rectification, while LPS-RS*U* (1 µg/ml) reversed AMPAR current rectification (Fig. 2e-f). Together, the data from cultured *WT* and *TLR4^-/-^* neurons demonstrate that neuronal TLR4 is contributes to the increase in CP-AMPAR currents *in vitro* and is sufficient to modulate CP-AMPAR currents.

**Fig. 2.**
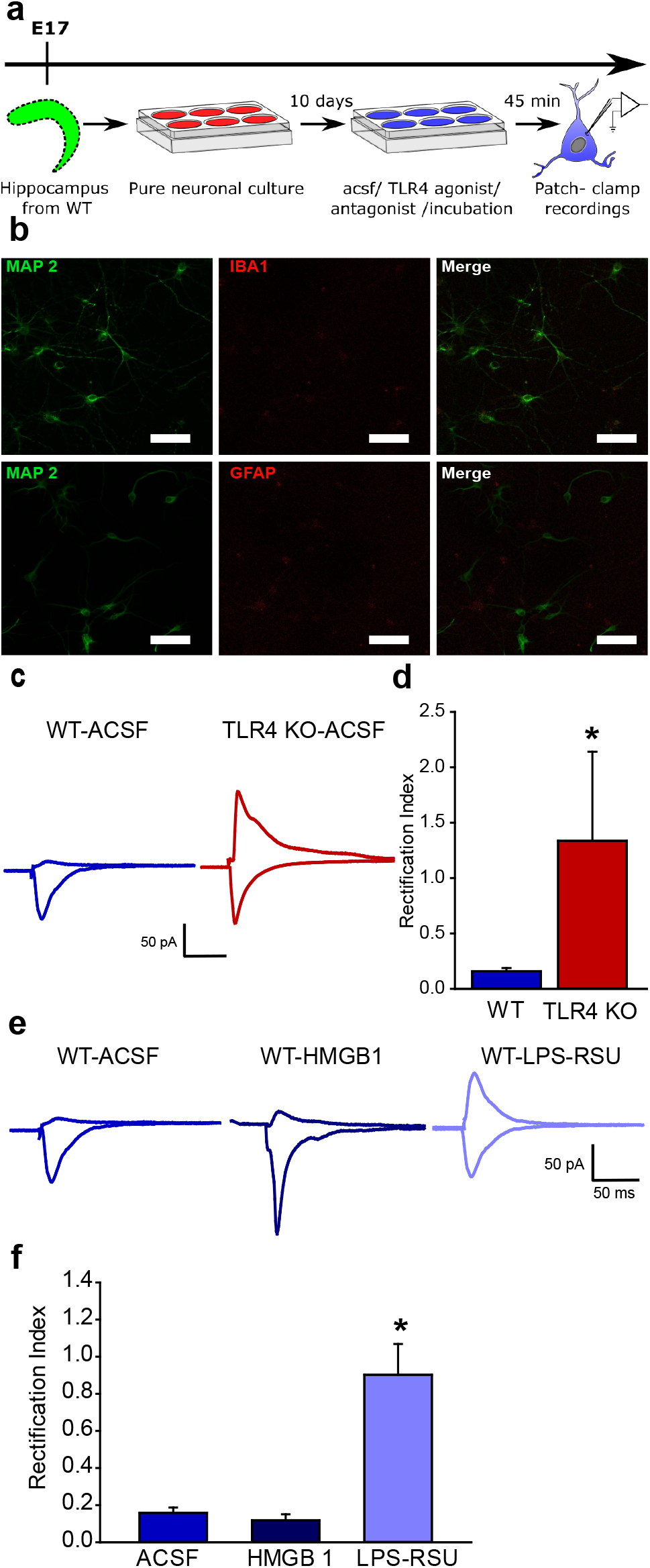
TLR4 modulation of calcium-permeable AMPAR currents in hippocampal neuron-only cultures *in vitro*. a. Schematic of the experimental paradigm illustrates the timeline for isolation of the hippocampus and maintenance in vitro followed by electrophysiology. b. Representative maximum intensity projection of confocal image stacks from a neuronal culture plated at embryonic day 17 and stained for MAP2 at DIV 12 to reveal neurites. c. Representative traces of AMPAR current recordings in neurons cultured from *WT* (left) and *TLR4^-/-^* mice (right) in response to a 1 mA stimulus to the neuropil. Recordings were obtained from holding potentials of −60 mV and +40 mV as in Fig. 1f. Note the AMPAR current rectification in neurons from *WT* and symmetric currents in *TLR4^-/-^*. d. Summary plot of rectification index measured as the ratio of peak AMPAR current at +40 mV to the peak current at −60 mV. (* indicates p<0.05 by Mann-Whitney test). e. Sample AMPAR current traces recorded in response to neuropil stimulation in *WT* neuronal cultures. Neurons were held at +40mV and −60 mV to obtain recordings in ACSF (left), TLR4 agonist HMGB1 (middle) and TLR4 antagonist LPS-RS*U* (right). Note the AMPAR current rectification in the absence of drugs and symmetric currents in LPS-RS*U*. f. Summary plot of rectification index. * indicates p<0.05 by One Way ANOVA on ranks followed by post-hoc Tukey’s test.

### Systemic TLR4 antagonism *in vivo* has opposing effects on early dentate excitability and epileptogenicity in control and injured brains

Next, we focused on the impact of the early post-injury TLR4 signaling on subsequent epileptogenesis which has been shown to occur after fluid percussion injury (38, 39). Latency to seizures evoked by low-dose kainic acid (KA, 5 mg/kg i.p.) challenge was used to assess epileptogenicity (29). While only a few sham animals injected with low-dose KA developed electrographic and behavioral seizures (Fig. 3a-b), rats one month after brain injury reliably developed seizures within 30 minutes of KA injection. Latency to KA-induced seizures was significantly reduced in rats one month after brain injury (Fig. 3c). Additionally, sham injured rats reached a maximum Racine score of 2 or 3 and brain injured rats reached maximum score of 5 (Fig. 3d). Overall, compared to sham-injured controls, brain-injured rats develop more-severe seizures with shorter latency following low-dose KA injection demonstrating post-traumatic enhancement of seizure susceptibility. In a small cohort, we monitored animals for spontaneous seizures after brain injury. In 30-40-hour simultaneous video- and surface EEG recordings from rats 12-15 weeks after sham or FPI, none of the sham rats (n=9) showed electrographic seizures, while spontaneous electrographic and behavioral seizures were observed in 8 of 13 FPI rats (∼62%, Fig. 4a-d). In rats with spontaneous seizures the average seizure severity was Racine score 3.37±0.18 (range of 3-4) and average seizure duration was 97.88±10.20 sec (range of 50-140 sec) confirming the risk for post-traumatic epileptogenesis in the experimental model.

**Figure 3.**
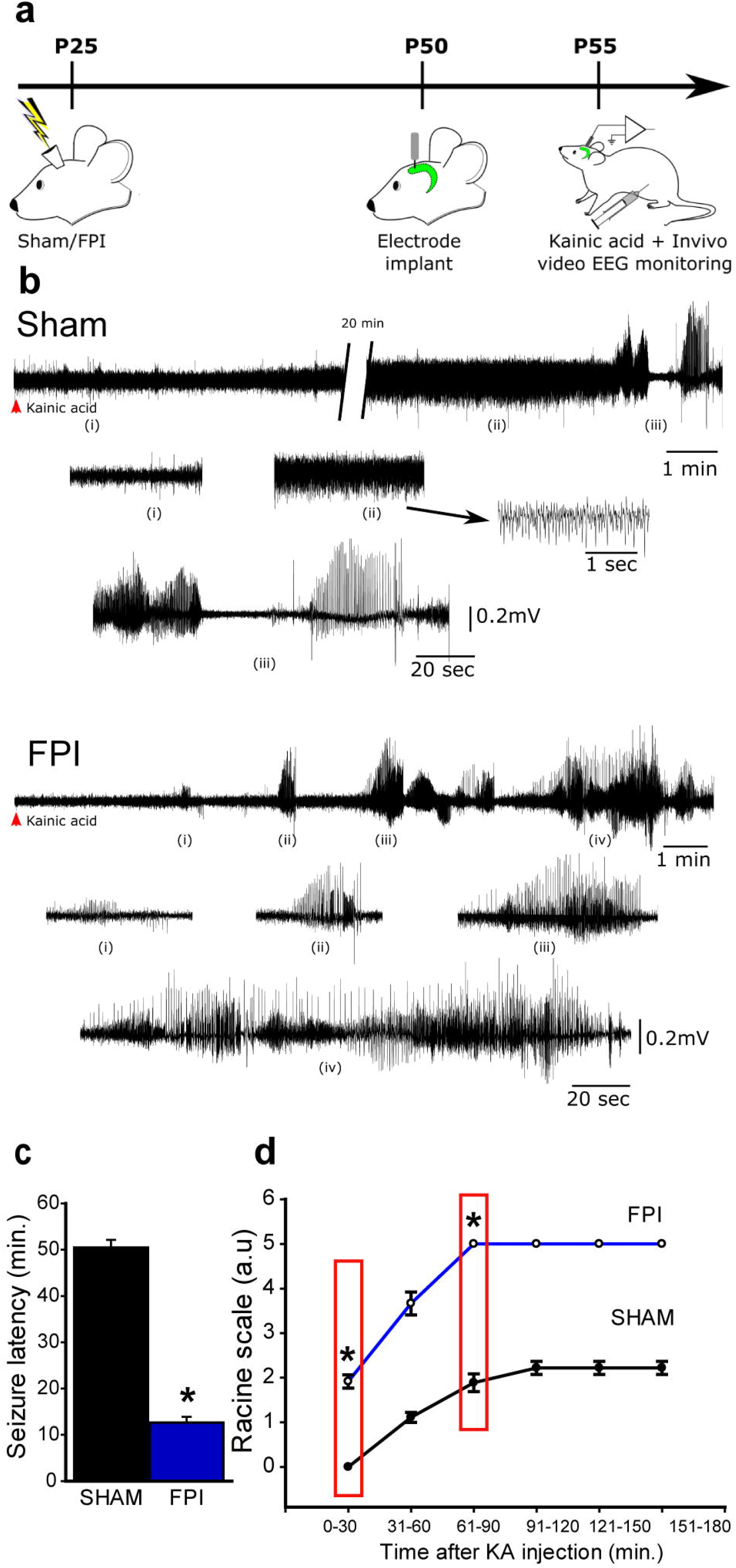
Enhanced susceptibility to chemically evoked seizures after brain injury. a. Schematic of timeline for *in vivo* injury followed by electrode implantation and low-does Kainic Acid challenge b. Sample hippocampal depth electrode recordings show evolution of electrographic activity following kainic acid injection in rats one month after sham (above) and FPI. Note that FPI animals develop electrographic seizures early which quickly develops into convulsive seizures (FPI, i-iv). c. Summary of latency to kainic acid induced seizures in rats 1-month after sham or brain injury. d. Summary of progression of seizure severity by Racine’s scale over time in sham (n=9) and FPI (n=12) rats. Boxed areas indicate time points at which statistical comparisons are reported. * indicates p<0.05 by Student’s t-test.

**Fig. 4.**
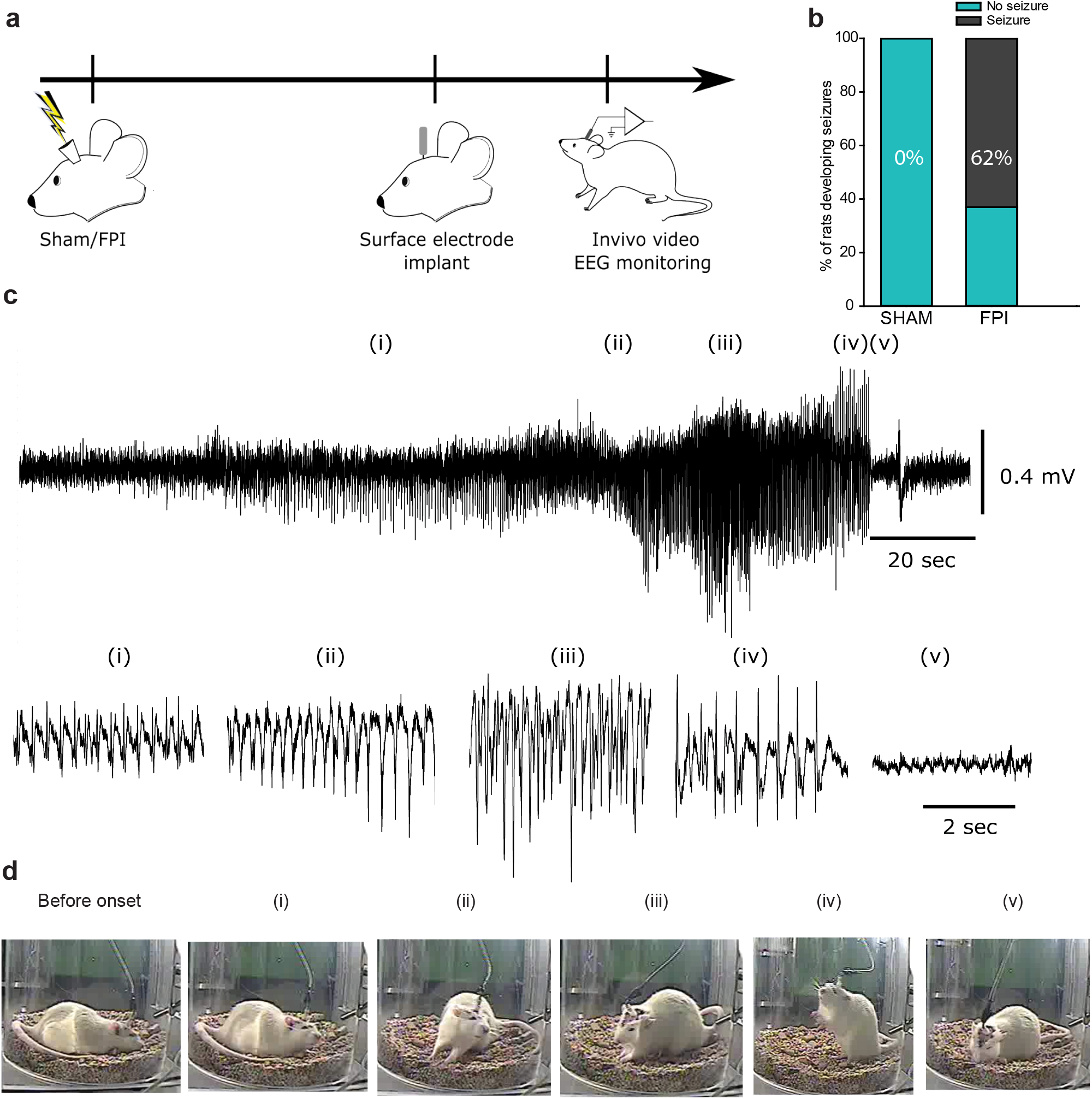
TLR4 antagonism alters delayed occurrence of spontaneous epileptic seizures. a. Timeline for assessment of spontaneous seizures following sham or FPI in rat. b. Summary plot illustrates percentage of rats developing spontaneous seizures. c. Sample subdural electrode recordings in brain injured rat obtained 12-15 weeks after injury. Panels below show expanded segments of EEG data from regions i-v. d. Video snapshots corresponding to segments i-v are illustrated. The time of recoding first ictal activity is noted. * indicates *p*<0.05 by z-test based on 10 sham and 13 FPI rats.

To transition our findings on TLR4 modulation of excitability in *ex vivo* slices to *in vivo*, we examined whether systemic TLR4 antagonism could modify dentate excitability one week after injury. Sham and FPI rats were treated with CLI-095 (0.5mg/kg, s.c. for 3 days) starting 20-24 hours after injury and dentate excitability was examined in hippocampal slices 6-8 days later. As with acute incubations, CLI-095 treatment *in vivo* increased dentate excitability in sham-injured rats, while effectively suppressing excitability after FPI (Fig. 5a-c). These data justify the use of *in vivo* treatments in evaluating the effect of TLR4 signaling on early dentate excitability and epileptogenesis after brain injury.

**Fig. 5.**
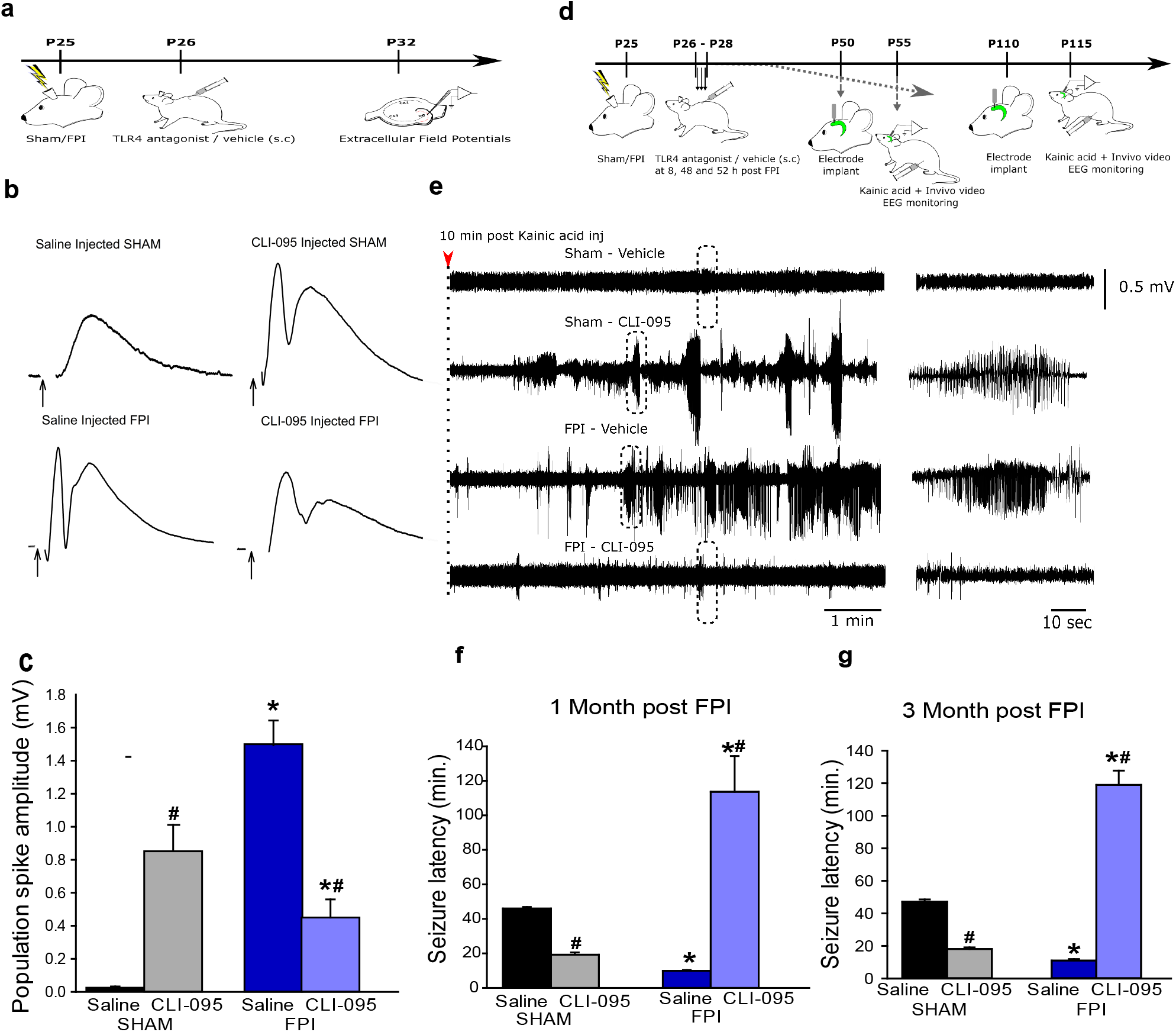
Systemic TLR4 antagonism *in vivo* modulates dentate network excitability in both sham- and brain-injured rats. a. Illustration of timeline for *in vivo* treatments and slice physiology in sham- and brain-injured rats. b. Dentate population responses evoked by a 4-mA stimulus to the perforant path in slices from sham (above) and brain injured (below) rats treated *in vivo* with saline (left) and CLI-095 (0.5mg/kg, one day after FPI, right). Arrows indicate truncated stimulus artifact. c. Summary data demonstrate the effect of CLI-095 on perforant path-evoked granule cell population spike amplitude in slices from sham (6 slices each from 3 rats) and FPI rats (6 slices each from 3 rats). Error bars indicate s.e.m. * indicates *p*<0.05 from sham and ^#^ indicates p<0.05 compared to saline within injury type, by TW-ANOVA followed by Tukey’s post hoc test. d. Illustration of timeline for surgical procedures, *in vivo* treatments and EEG recordings in sham and brain injured rats. e. Sample hippocampal depth electrode recordings in sham and brain injured rats treated with vehicle or CLI-095 (0.5mg/kg). Recordings show EEG activity 10 minutes after KA injection (right panel). Expanded traces of the activity in boxed areas illustrated in the left panels show lack of seizure activity in sham-vehicle and FPI-drug conditions and development of seizure activity in FPI-vehicle and sham-drug rats in the 10-20-minute period after KA injection. f-g. Summary of latency to KA-induced seizures in rats 1-month (f) and 3-months (g) after sham or brain injury. * indicates *p*<0.05 from sham and ^#^ indicates p<0.05 compared to saline within injury type, by TW-ANOVA followed by Tukey’s post hoc test.

Although the post-injury increase in dentate excitability is postulated to underlie a long-term increased risk for epilepsy (4, 5, 40), this association remains untested. Since blocking TLR4 reduces excitability after injury and increases excitability in uninjured rats, TLR4 antagonism provides an opportunity to test the link between early excitability and subsequent risk for seizures. Rats were treated with systemic CLI-095 (0.5mg/kg) or saline once daily for 3 days starting 24 hours after sham or brain injury, when hippocampal TLR4 levels were found to be maximal (12, 41). As expected, saline-treated injured rats had a shorter latency to KA-induced seizures compared to saline-treated sham rats when examined 1 and 3 months after injury (Fig. 5d-f). CLI-095 treatment decreased latency to KA-evoked seizures in sham injured rats while prolonging seizure latency in FPI rats at one-month and three months post-FPI (Fig. 5f-g). Mechanistically, the data reveal that the risk for seizures is driven by early changes in dentate excitability. Translationally, these results show that transiently blocking TLR4 early after brain injury has the potential to prevent post-traumatic epileptogenesis.

### Early focal TLR4 antagonism is sufficient reduce post-traumatic increases in seizure susceptibility while delayed treatment is ineffective

To probe the association between hippocampal TLR4 signaling and epileptogenesis, we adopted a focal TLR4 antagonist treatment. Ipsilateral bolus hippocampal injection of LPS-RS*U* (2mg/ml) 24 hours after FPI/sham led to decreased latency to KA-evoked seizures in sham rats and eliminated seizures altogether in FPI rats when examined one and three months after brain injury (Fig. 6a, b, d). Hippocampal LPS-RS*U* treatment enhanced seizure severity in controls and reduced seizure severity in FPI rats at one and three months after FPI/sham. (Fig. 6c, e). Thus, local suppression of TLR4 in the injured hippocampus and its modification of early dentate excitability can predict long-term seizure susceptibility.

**Fig. 6.**
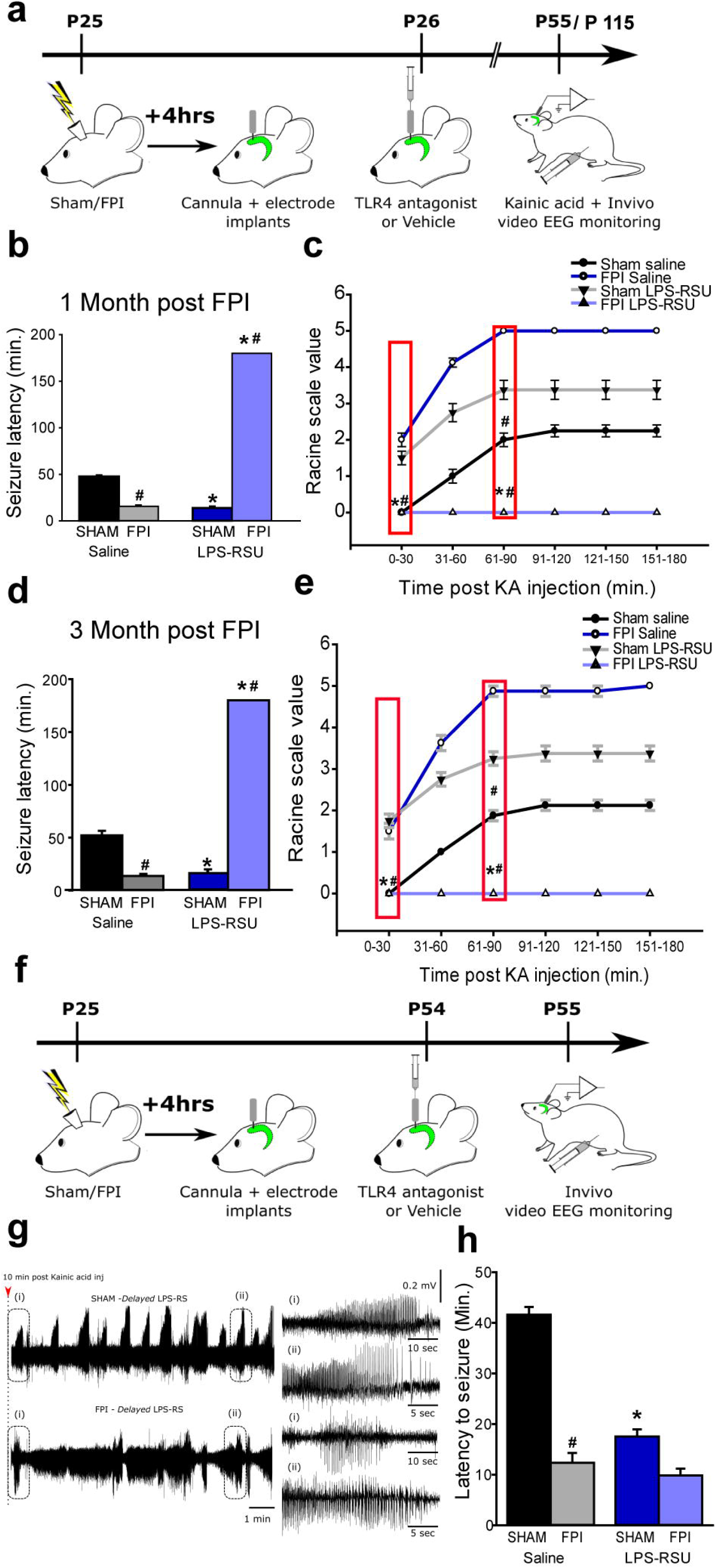
Early but not delayed focal hippocampal TLR4 antagonism eliminates post-traumatic increase in seizure susceptibility. a. Schematic of treatments and surgical procedures relevant to panels b-e for experimental rats. b and d. Summary of latency to KA (5 mg/kg) induced seizures in sham and FPI rats 1-month (b) and 3-months (d) post injury obtained using hippocampal depth electrodes. Rats were injected with saline or LPS-RS*U* (2mg/ml, single bolus injection) unilaterally on the side of the FPI implant 24 hours after injury. c and e. Progression of seizure severity and maximum seizure score by Racine’s scale reached in saline or drug treated sham and FPI rats 1-month (c) and 3-months (e) post injury. * indicates *p*<0.05 from sham and ^#^ indicates p<0.05 compared to saline within injury type. N=8 rats per group. f. Illustration of timeline for delayed treatment. g. Sample hippocampal depth electrode recordings in sham and brain injured rats treated with LPS-RS*U* (2mg/ml) one-month post FPI. EEG recordings show rapid development of seizures within 10-15 minutes after KA injection. Panels to the left show expanded traces from the boxed areas. h. Summary plot of latency to kainic acid induced seizures. * indicates *p*<0.05 from sham and ^#^ indicates p<0.05 compared to saline within injury type, by TW-ANOVA followed by Tukey’s post hoc test.

Interestingly, analysis of mortality among experimental animals revealed that sham-injured rats treated with TLR4 antagonists had increased mortality while post-traumatic rats treated with TLR4 antagonists showed lower mortality (% mortality between one week after treatment and experimental end point: sham-saline: 0%, 0/20; Sham-LPS-RS*U*: 26.9%, 7/26; FPI-saline: 44.4%, 16/36; FPI-LPS-RS*U*: 13.3%, 4/30). Moreover, evaluation of the spontaneous seizures in TLR4 antagonist-treated sham and FPI animals revealed that one LPS-RS*U*-treated sham rat (of 6 examined; 16.7%) and none of the saline-treated sham rats developed spontaneous seizures. However, three (of 6 tested; 50%) saline-treated FPI rats exhibited spontaneous seizures and none of the 6 LPS-RS*U*-treated FPI rats showed electrographic or behavioral seizures in the same duration of testing.

The salience of the early period of excitability to seizure susceptibility, was tested by delivering focal LPS-RS*U* or saline treatments 1 month after injury followed by evaluation of KA-induced seizures on the following day. Delayed LPS-RS*U* treatment reduced the latency to seizures in uninjured rats confirming that blocking TLR4 can modify network excitability in controls (Fig. 6f-h). Conversely, delayed LPS-RS*U* treatment failed to reduce seizure susceptibility after FPI (Fig. 6f-h), highlighting the importance of the early post-injury increases in dentate network excitability in long-term seizure risk.

### TLR4 antagonism reduces cellular inflammation following FPI without altering inflammatory milieu in controls

Since TLR4 signaling leads to cellular inflammation, prior studies have attributed beneficial effects of blocking TLR4 after brain trauma to reducing inflammatory responses (42). To test whether inflammatory responses could mediate the divergent effects of TLR4 antagonists on seizure susceptibility, we examined the inflammatory responses in hippocampal tissue obtained from saline/CLI-095-treated rats three days after sham/FPI. In western blots, CLI-095 significantly reduced hippocampal expression of TLR4, GFAP and Iba1 in the injured brain without enhancing expression in sham animals (Fig. 7a-e). A cohort of animals were examined one month after FPI and saline/CLI-095 treatment. Consistent with our previous findings demonstrating that TLR4 returns to control levels by one-month (12), hippocampal expression of TLR4 expression in saline treated FPI rats was not different from sham-saline rats when examined one month after treatment (Sham-saline: 1.00±0.0; FPI-Saline: 1.10±0.28, p=0.92 by Tukey’s post hoc test, n= 3 each). TLR4 expression in FPI rats treated with CLI-095 one day after injury was not different from sham-saline or FPI-saline groups (TLR4 expression normalized to sham-saline: FPI-CLI 0.82±0.18, p=0.58 by One-Way ANOVA, Tukey’s post hoc test, n= 3 each). Curiously, GFAP protein levels were highly variable after FPI, increasing in some and decreasing in others and was not significantly different between groups (FPI-saline normalized to sham Saline, Sham-saline: 1.00±0.0, n=3; FPI-Saline: 1.74±0.67, p=0.35, n=4). However, GFAP protein levels in FPI-CLI group trended to decrease in FPI-saline group one-month post-injury (GFAP expression in FPI-CLI normalized to sham-saline, FPI-saline: 1.74±0.67, n=4; FPI-CLI: 0.17±0.06, p=0.063, n=4 by Tukey’s post hoc test). Finally, we examined whether FPI and early CLI-095 treatment altered GluA1 expression one-month after injury. Surface expression of GluA1 was not different between saline treated sham and FPI groups (surface/total GluA1 ratio normalized to sham-saline: sham-saline: 1.00±0.0; FPI-Saline: 1.12±0.29, p=0.94, n=3 each). Additionally, early CLI-095 treatment did not alter surface-expression of GluA1 tested one month after FPI (surface/total GluA1 ratio normalized to sham-saline: FPI-CLI 1.23±0.43, p=0.96, n=3 each). However, total GluA1 levels showed a significant increase one month after FPI (total GluA1 normalized to sham-saline, sham-saline: 1.00±0.0; FPI-Saline: 1.39±0.15, p=0.03, n=4 each) and reduced significantly following early CLI treatment (total GluA1 normalized to sham-Saline: FPI-Saline: 1.39±0.15, FPI-CLI: 0.75±0.05, p=0.002, n=4 each, by One-Way ANOVA, Tukey’s post hoc test). These changes in total GluA1 expression are consistent with persistent post-FPI increase in hippocampal excitability after FPI (43) and demonstrates that early treatment with CLI-095 reduces a potential molecular substrate for persistent posttraumatic increase in excitability.

**Fig. 7.**
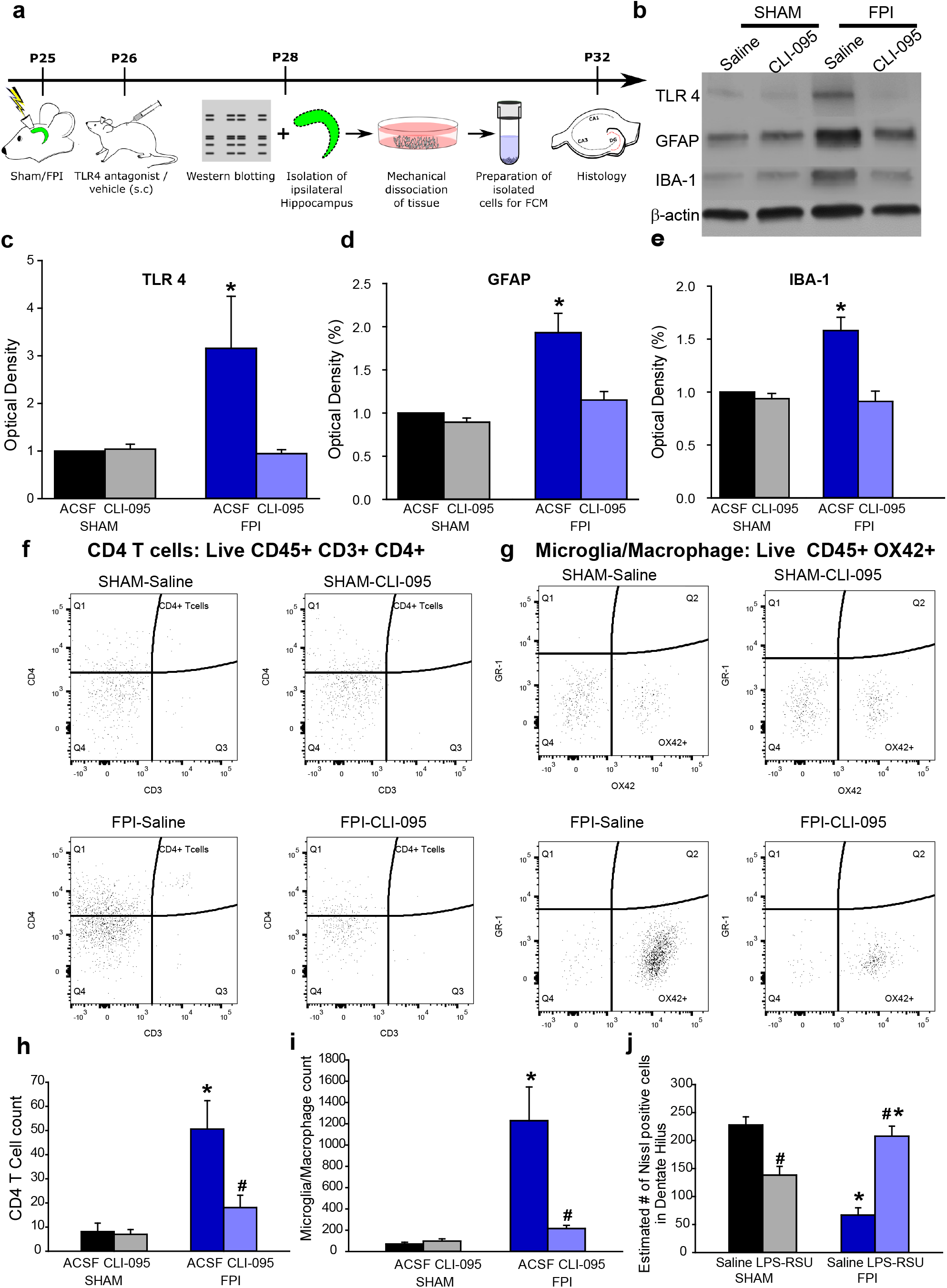
TLR4 antagonism suppresses cellular inflammation and loss after brain injury without perturbing inflammatory response in uninjured sham rats. a. Experimental timeline for western blot, flow cytometry and histological studies. b. Representative western blots for TLR4, GFAP and IBA-1 in hippocampal samples from the injured side obtained 3 days after saline/drug treatment. Treatments began 24 hours after injury. Corresponding β illustrated. c-e. Summary histograms of expression of TLR4 (c), GFAP (d) and IBA-1 (e) as a % of the expression levels in sham-saline treated controls. f. Representative CD4/CD3 scatter plots from the hippocampus on the injured side of saline- and (left) and CLI-095 treated (right) sham (above) and brain injured (below) rats. CD4/CD3 scatter plots were gated on live CD45+ cells. The population of interest is noted by the red oval. g. Representative GR-1/Ox42 scatter plots from the hippocampus on the injured side of saline- and (left) and CLI-095 treated (right) sham (above) and brain injured (below) rats. GR-1/OX42 scatter plots were gated on live CD45+ cells. h-i. Quantification of total CD45^+^CD3^+^CD4^+^ T cells (i) and CD45^+^OX42^+^ microglia in the experimental groups. Data are presented as mean ± s.e.m., n = 12 animals/treatment (4/group with 3 replicates) * indicates *p*<0.05 from sham and ^#^ indicates p<0.05 compared to saline within injury type by Kruskal-Wallis One Way ANOVA on ranks followed by post-hoc pairwise multiple comparison by Student-Newman-Keuls Method. j. Estimate of Nissl stained cells/ section in the ipsilateral dentate hilus in the experimental groups. n = 11 slices in 3 rats in sham-saline, 11 slices in 3 rats for sham treatment 12 slices in 4 rats from FPI-saline and 11 slices from 4 rats in FPI-treatment. * indicates *p*<0.05 from sham and ^#^ indicates p<0.05 compared to saline within injury type by TW-ANOVA followed by post-hoc Tukey’s test.

Flow cytometry to quantify the cellular inflammation in the hippocampus was performed three days after sham/FPI followed by treatments. The data revealed a significant increase in CD45^+^CD3^+^CD4^+^ T-cells and CD45^+^Ox42^+^ microglia/macrophages in saline-treated rats compared to uninjured controls consistent with timeline for post-injury inflammatory responses (44). CLI-095 treatment reduced T-cell and microglial/macrophage count in the hippocampus of brain injured rats (Fig. 7f-i). Contrary to its effects on excitability, CLI-095 failed to increase T-cells or microglia/macrophage counts in sham rats (Fig. 7f-i). To directly assess the effect of treatment on histopathology, we examined the dentate hilus for neuronal loss one week after injury and treatments. Nissl staining for surviving hilar neurons demonstrated that TLR4 antagonists (ipsilateral intrahippocampal LPS-RS*U*) prevented hilar cell loss observed after brain injury and reduced hilar cell counts in uninjured rats (Fig. 7j) demonstrating that early cell loss tracks changes in excitability regardless of the inflammatory milieu in controls.

### Driving excitability limits ability of TLR4 antagonists to suppress seizure susceptibility after brain injury

To test whether the ability of TLR4 antagonists to suppress excitability can influence post-injury seizure susceptibility independently of its ability to reduce inflammation we tested whether enhancing AMPAR currents during TLR4-antagonism would increase excitability downstream of TLR4. Ampakines, positive allosteric modulators of AMPARs, have been used *in vivo* to enhance AMPAR currents (45, 46). Since TLR4 modulates AMPAR currents only after brain injury, we first validated the ability of an Ampakine to augment AMPAR currents in the presence of TLR4 antagonists in slices from brain-injured rats. Afferent-evoked granule cell AMPAR current charge transfer was enhanced one week after brain injury (Fig.8a-c) and reduced by incubation in LPS-RS*U* (Fig. 8d-e). Incubation in LPS-RS*U* (1µg/ml) together with an Ampakine (CX546; 300µM) enhanced charge transfer confirming that this Ampakine can enhance AMPAR currents in the presence of LPS-RS*U* (Fig. 8d-e). Following *ex vivo* validation, rats one day after FPI were treated with LPS-RS*U* followed by an Ampakine (CX546; 300µM) delivered through an implanted hippocampal cannula electrode (Fig. 8f). Although local treatment with LPS-RS*U* early after injury eliminated seizures during the KA-challenge, simultaneous administration of an Ampakine resulted in development of seizures and an apparent reduction in seizure latency demonstrating a role for TLR4-mediated excitability in seizure susceptibility. However, the latency to seizures in injured animals receiving combined LPS-RS*U* and Ampakine treatment was longer than those treated with saline (Fig. 8f-h). These results identify TLR4-mediated enhancement of glutamatergic currents after brain injury as critical contributor to post-traumatic epileptogenesis.

**Fig. 8.**
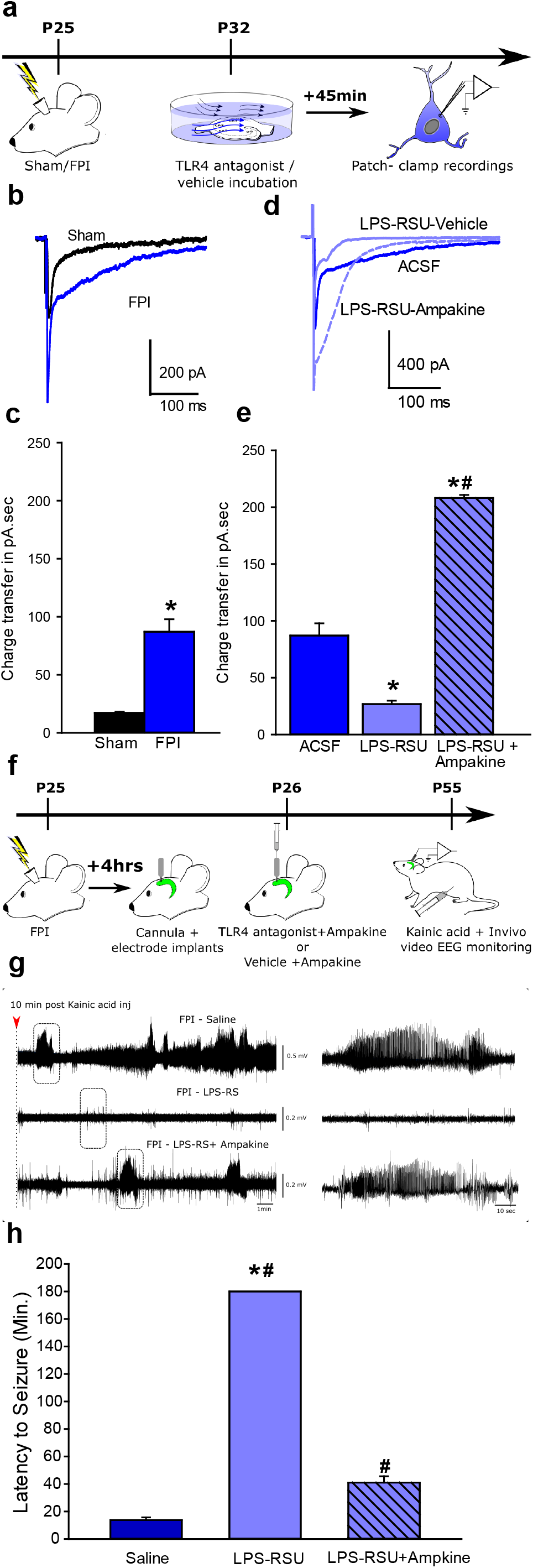
Augmenting AMPAR currents increases seizure susceptibility in TLR4 antagonist-treated brain-injured rats. a. Schematic for experiments testing the efficacy of Ampakine to enhance perforant path-evoked AMPAR currents in *ex vivo* preparations. b. Overlay of afferent evoked AMPAR currents in slices from sham (black) and FPI (blue traces) rats. c. Summary histograms of AMPAR charge transfer in experimental groups. *indicate p<0.05 by Mann Whitney test. d. Overlay of afferent-evoked AMPAR currents in slices from FPI rats incubated in ACSF/saline (dark blue), LPS-RS*U* and saline (light blue solid line) and LPS-RS*U* and CX546 (light blue dashed line). e. Summary histograms of AMPAR charge transfer in experimental groups. *indicate p<0.05 by One Way ANOVA on ranks followed by post-hoc Tukey’s test for differences between slices from FPI following different drug incubations. The same data set for FPI-ACSF is used in panels c and e. f. Illustration of treatments and timeline for testing the ability of CX546 to alter seizure susceptibility. g. Sample hippocampal depth electrode recordings in brain injured rats treated with saline or LPS-RS*U* (2mg/ml) or LPS-RS*U* (2mg/ml) + CX546 (300µM). EEG recordings obtained 10 minutes after KA (5 mg/kg, i.p) injection shows the early development of seizures in saline treated FPI rats (above), lack of seizure activity in FPI rats treated with LPS-RS*U* 24 hours after injury (middle) and development of seizure activity in FPI rats treated with CX546and LPS-RS*U* (lower). Expanded traces in the boxed areas are shown (right panels). h. Summary plot of latency to KA induced seizures. Saline and LPS-RS*U* data in h were presented in 5b. * indicates *p*<0.05 from saline and ^#^ indicates p<0.05 compared to LPS-RS*U* by one-way ANOVA followed by Tukey’s post hoc test.

## Discussion

This study reveals a Janus head-like dichotomy in the effect of TLR4 signaling in vivo on neuronal excitability in the uninjured versus injured hippocampal dentate gyrus, which impacts network excitability and seizure susceptibility. Our results highlight the complex interactions between immune signaling, brain injury, and epileptogenesis at distinct functional levels and temporal scales. At a time scale of minutes to days, TLR4 signaling enhanced GluA1 surface expression and CP-AMPAR currents in the injured brain which led to enhanced network excitability. TLR4 antagonist treatment, *ex vivo* and *in vivo*, reversed post-traumatic increases in TLR4 and GluA1 surface expression and network excitability to control levels (Figs. 1 and 5). Notably, TLR4 modulation of CP-AMPAR currents was observed in isolated neuronal cultures, revealing a direct effect of neuronal TLR4 in regulating excitability, independent of glial contribution. It is attractive to speculate that TLR4 recruits glia-independent signals to promote GluA1 trafficking and surface expression in the injured brain. These results constitute the first demonstration of a functional role for central neuronal TLR4 in modulating excitability and AMPAR currents.

In contrast to its effects after injury, TLR4 antagonists enhanced network excitability in controls (12). in contrast to effects on excitability, TLR4 antagonists suppressed the post-traumatic cellular inflammatory responses without inducing inflammation in controls. Although the mechanisms by which TLR4 signaling increases dentate excitability in controls is currently unknown, our findings demonstrate that enhanced inflammatory responses do not underlie these effects. Moreover, consistent with our prior study (12), TLR4 does not modulate glutamate currents in controls. Of note, our treatments included two mechanistically distinct antagonists, LPS-RS*U* delivered locally and CLI-095 administered systemically, to demonstrate the divergent effects of TLR4 signaling on excitability and seizure susceptibility in controls and after injury. These findings suggest that a distinct TLR4 signaling pathway is utilized after injury. Overall, analysis of the short-term effects of TLR4 signaling revealed constitutive suppression of excitability under basal conditions and enhanced excitability mediated by neuronal CP-AMPAR currents after injury.

Our data demonstrating that transient perturbations in hippocampal TLR4 signaling have long-term implications for network excitability and stability constitutes a significant advance in understanding the relevance of TLR4 signaling for post-traumatic neuropathology since our earlier acute studies in vitro (12). Brain injury and TLR4 antagonism both affect TLR4 signaling, augment network excitability, and lead to increased susceptibility to seizures long-term. Blocking TLR4 signaling *in vivo* early *after* brain injury reduced network excitability and cellular inflammatory responses, hippocampal GluA1 expression and diminished seizure susceptibility in the long-term. However, transiently increasing AMPAR currents in the injured hippocampus reduced the ability of LPS-RS*U* to prevent chemically evoked seizures. These data show that, independent of the effect on inflammation, TLR regulation of excitability impacts seizure susceptibility. Moreover, transiently antagonizing TLR4 enhanced dentate excitability and promoted seizures without inducing cellular inflammation in controls. Therefore, regardless of effects on the inflammatory milieu, manipulations that increase in hippocampal dentate excitability drive long-term increased risk for epilepsy. Importantly, the aberrant neuronal TLR4-mediated increase in CP-AMPAR may be targeted *after* brain trauma to prevent epileptogenesis.

There is growing evidence for interactions between inflammatory signaling and neuronal physiology in the normal brain and in disease (6, 47–49). Brain injury and seizures lead to increases in endogenous damage associated molecular patterns (DAMPs), such as HMGB1, which can activate pattern-recognition receptors, including TLR4 (11, 41, 50, 51). While mice lacking TLR4 show improved neurological outcomes after brain injury, the beneficial effects have been attributed to reduced inflammation (52). TLR4 antagonists have been shown to reduce chemoconvulsive seizures by modulating NMDAR currents, suggesting that TLR4 modifies neuronal excitability (11, 13). Although TLR4 is expressed in neurons (11, 12), TLR4 modulation of NMDAR-dependent calcium entry was found to involve astroglial purinergic signaling or generation of the pro-inflammatory cytokine Tumor Necrosis Factor α (TNFα) (11, 13, 49, 53, 54). Accordingly, glial signaling has been proposed to underlie the neurophysiological effects of TLR4 (6, 49, 55). Our demonstration that TLR4 activity modulates neuronal CP-AMPAR currents, independent of glia, identifies a novel mechanism by which TLR4 promotes excitotoxic damage and epileptogenesis independent of inflammatory signaling. Our findings uncover a unique neuronal axis of TLR4 signaling similar to reports in Major Histocompatibility class I molecules (56). These data raise the possibility that neurons may repurpose the complexity of immune receptors to modulate neuronal excitability. Like enhanced excitability, cellular inflammatory response is a hallmark of traumatic brain injury (3, 57). While TLR4 blockers limited hippocampal cellular inflammation in brain injured animals and failed to induce inflammation in controls, they had an opposite effect on excitability and epileptogenicity depending on the injury profile. Augmenting AMPAR currents with an Ampakine alongside LPS-RS*U*, 24 hours after brain injury, decreased latency to seizures compared to LPS-RS*U* administered alone demonstrating that TLR4 enhancement of AMPAR currents contributes to epileptogenesis. However, latency to seizure in injured rats receiving an Ampakine and LPS-RS*U* was longer than following saline treatment, which may underscore a parallel beneficial effect of reducing inflammation. Taken together, our findings identify a neuronal axis for TLR4 signaling that enhances excitability after brain injury and demonstrate that suppression of both excitability and inflammatory responses contribute to anti-epileptogenic effects of TLR4 antagonists.

The contribution of glial mediators to TLR4 regulation of neuronal excitability has been identified in earlier studies (10, 13, 49). Our findings in neuronal cultures demonstrate that glial signals are not engaged in the TLR4 enhancement of CP-AMPAR currents, revealing neuronal TLR4 signaling in the CNS. Future studies are needed to address the molecular underpinnings of TLR4 signaling in CNS neurons as has been examined in peripheral neurons (16, 58, 59). We previously reported a ∼40% increase in granule cell peak AMPAR currents in slices from brain injured rats, which was abolished by incubation in TLR4 blockers (12). Our finding that blocking TLR4 in slices selectively decreases GluA1 surface expression after brain injury without altering GluA2 expression indicates that an increase in GluA2 lacking AMPAR with calcium permeable GluA1 containing receptors underlies the post-injury increase in AMPAR currents. This could occur from exocytosis and surface expression of a reserve pool of GluA1, similar to the processes mediated by glia-derived TNFα in culture systems (60). Increases in granule cell CP-AMPARs have been shown to contribute to excitotoxic damage after experimental stroke (34), suggesting that an increase in CP-AMPARs may drive excitotoxic neuronal loss and epileptogenesis after brain injury. Similarly, TLR4 also augments AMPAR currents in mossy cells after FPI (12) and enhanced mossy cell activity drives an increase in granule cell AMPAR charge transfer after brain injury (33). Thus, the rapid TLR4 regulation of AMPAR currents likely plays an important role in increasing excitability shortly after brain injury. In contrast to the lack of increase in total GluA1 levels one week after FPI, hippocampal total GluA1 expression appears increased one month after FPI suggesting that the mechanisms underlying enhanced excitability evolve with disease progression. However, CLI-095 treatment early after FPI significantly lowered GluA1 expression one month after injury demonstrating that the early reduction in excitability limits long term increases in GluA1 expression and seizure susceptibility.

We find that blocking TLR4 signaling in uninjured rats promotes neuronal loss in the dentate hilus and raises the concern that systemic TLR4 antagonists may contribute to cell loss in uninjured brain regions and compromise circuit function. These findings underscore the need to resolve the distinct mechanisms underlying TLR4 regulation of neuronal excitability in the injured and uninjured brain to enable selective targeting of the pathological mechanisms. Curiously, AMPAR current rectification in cultured neurons from *WT* mice resembled the injured brain while those from *TLR4^-/-^* mice were similar to the uninjured brain. This demonstration that TLR4 is necessary for AMPAR current rectification in cultured neurons provides a mechanism for the previously reported developmental increase in CP-AMPARs in neuronal cultures (36). The parallels between TLR4 effects on AMPAR currents in cultured neurons and the injured brain suggest that the process of generating cultures may impart an “injured” AMPAR phenotype.

Although early increases in dentate excitability after brain injury have been proposed to contribute to epileptogenesis (4–6), a convincing demonstration of a causal association has been lacking. We recently showed that suppressing post-traumatic increases in neurogenesis reduced dentate excitability and epileptogenesis in parallel, supporting the association between excitability and epileptogenesis (18). By identifying that early excitability, regardless of injury phenotype or inflammatory milieu, predicts epileptogenesis, our study supports the causal association between enhanced dentate excitability and remote development of epilepsy. Indeed, our experiments using two TLR4 antagonists, LPS-RS*U*, an extracellular blocker, and CLI-095, a blocker of the intracellular signaling domain (32), demonstrated the same paradoxical effects on excitability in control and brain-injured animals and eliminate the possibility of nonspecific effects. Moreover, CLI-095 was administered systemically while LPS-RS*U* was administered by focal injection, demonstrating that both systemic and focal TLR4 antagonism could enhance epileptogenesis in controls and reduce epileptogenesis after brain injury. Interestingly, a recent study found that blocking HMGB1 prior to pediatric brain injury in mice failed to reduce seizure susceptibility (61). It is possible that suppression of HMGB1 prior to injury rather than after injury, and recruitment of TLR4 signaling by alternative injury-induced DAMPs may have contributed to the inability of HMGB1 to block seizures. While early TLR4 antagonist treatment after injury limits seizure susceptibility, delayed treatment is ineffective in reducing seizure susceptibility (Fig. 6d-f), validating the critical role for early network excitability and a potential “therapeutic window” in post-traumatic epileptogenesis.

Traumatic brain injury is a major risk factor for epilepsy among young adults. Consistent with earlier studies (39, 62), 50-60% of rats develop spontaneous seizures 12-15 weeks after moderate FPI. It is possible that strain differences, use of younger juvenile rats which are more susceptible to adverse effects of brain injury (63), implementation of injury on the day after surgery to minimize neuroprotection due to surgical anesthesia and inclusion criteria based on apnea duration contributed to greater proportion of rats developing spontaneous seizures compared to earlier studies (64–66). The ability of TLR4 antagonists to reduce seizure susceptibility when administered 24 hours *after* brain injury presents an opportunity to target TLR4 to prevent post-traumatic epilepsy. Indeed, CLI-095 (Resatorvid) crosses the blood-brain-barrier, has a half-life of 6 hours (20), and is FDA approved for clinical trials (67). However, its proepileptogenic effect and potential for inducing neuronal loss in uninjured animals poses a major impediment to systemic use of TLR4 antagonists. Since we identify that TLR4 regulation of CP-AMPARs is unique to the injured neurons, elucidating the specific molecular pathways underlying TLR4 enhancement of AMPAR currents after injury would enable targeted suppression of the pathological mechanisms.

## Acknowledgements

We thank Drs. Rizie Kumar and Amy Davidow for support with statistical analysis of flow cytometry data, Dr. Roman Shirakov for help with culture studies and Dr. Kelly Hamilton for helpful comments and discussions. This research was supported by CURE Foundation CF 259051, NJCBIR CBIR14RG024, NIH/NINDS R01 NS069861 and R01NS097750 to V.S. and NJCBIR CBIR15FEL011 to A.K.

## Author Contributions

Conception and Study Design: A.A.K, S.E., and V.S; Acquisition and analysis of data: A.A.K, Y.L., Di.S., De. S., S.S., J.G, B.S, K.K and A.P; Interpretation of results and preparation of figures: A.A.K, Di.S, S.S., and De.S. and S.E; Drafting manuscript and figures: A.A.K., De.S. and V.S.

## Potential Conflicts of Interest

Nothing to report

---

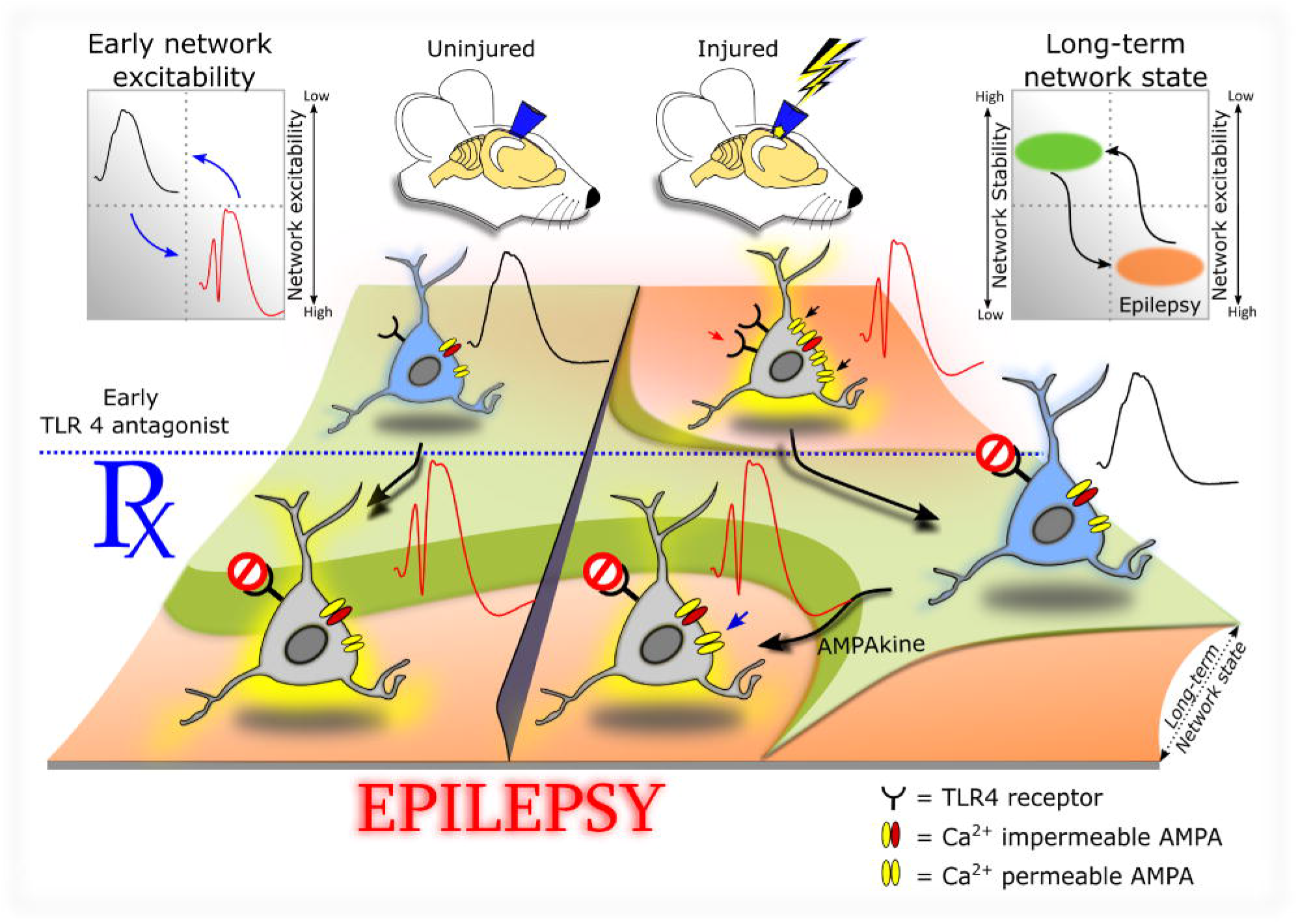

## Notes

#### Summary of Updates

Figure 1 and discussion section updated

