## supplemental data tables for "TLR4 signaling in neurons enhances calcium-permeable AMPAR currents and drives post-traumatic epileptogenesis"

**Table 1: Data as mean ± sem**

| **Fig #** | **units** | **Sham-ACSF/Saline** | **Sham Drug (CLI095 or LPS-RSU)** | **FPI**  **ACSF/saline** | **Sham Drug (CLI095 or LPS-RSU)** |
| --- | --- | --- | --- | --- | --- |
| 1d | mV | 0.21±0.08 | 1.04±0.17 | 1.77±0.49 | 0.14±0.11 |
| 1e | mV/ms | 0.96±0.13 | 1.99±0.15 | 2.52±0.34 | 0.83±0.18 |
| 1g | a.u | 0.90±0.05 | 1.03±0.12 | 0.12±0.02 | 0.67±0.06 |
| 1i | a.u | 100±0.0 | 101.7±4.56 | 137.18±8.4 | 110.18±5.34 |
| 1j | a.u | 100±0.0 | 105.34±7.30 | 108.96±9.67 | 103.78±7.25 |
| 1k | a.u | 100±0.0 | 108.80±5.50 | 108.8±6.96 | 108.00±7.74 |
| 1l | a.u | 100±0.0 | 107.80±3.83 | 106.00±6.54 | 112.10±8.50 |
| 5c | mV | 0.02±0.01 | 0.85±0.16 | 1.50±0.15 | 0.45±0.11 |
| 5f | min. | 46±0.89 | 19.22±1.34 | 9.85±0.44 | 113.66±20.71 |
| 5g | min. | 47.12±1.44 | 18.12±1.00 | 11±0.96 | 119.00±8.70 |
| 6b | min. | 47.71±1.76 | 15.62±1.54 | 13.85±2.12 | 180±0 |
| 6d | Min. | 52.15±4.27 | 15.63±1.5 | 13.75±1.96 | 180±0 |
| 6h | min. | 41.62±1.53 | 17.57±1.39 | 12.37±1.91 | 9.85±1.35 |
| 7c | a.u | 1.00±0 | 1.04±0.10 | 3.16±1.09 | 0.95±0.08 |
| 7d | a.u | 1.00±0 | 0.89±0.04 | 1.93±0.22 | 1.15±0.09 |
| 7e | a.u | 1.00±0 | 0.93±0.50 | 1.58±0.12 | 0.91±0.09 |
| 7h | # of cells | 8.17±3.47 | 7.01±1.95 | 50.60±11.75 | 18.09±5.10 |
| 7i | # of cells | 70.17±16.49 | 97.78±20.68 | 1228.93±316.49 | 215.60±29.91 |
| 7j | # of cells | 227.81±14.55 | 138.33±15.59 | 66.80±13.07 | 207.77±17.95 |
| **Fig #** |  | **WT** | **KO** |  |  |
| 2d | a.u | 0.16±0.02 | 1.33±0.80 |  |  |
| **Fig #** |  | **ACSF** | **HMGB1** | **LPS-RSU** |  |
| 2f | a.u | 0.16±0.02 | 0.12±0.03 | 0.92±0.16 |  |
| **Fig #** |  | **SHAM** | **FPI** |  |  |
| 3c | min. | 50.44±1.71 | 12.63±1.24 |  |  |
| 4b | a.u | 0% | 62% |  |  |
| 8c | pA.sec | 17.21±0.93 | 87.02±10.83 |  |  |
| **Fig #** |  | **ACSF** | **LPS-RSU** | **LPS-RSU +Ampakine** |  |
| 8e | pA.sec | 87.02±10.83 | 26.73±3.02 | 208.95±2.71 |  |
| **Fig #** |  | **Saline** | **LPS-RSU** | **LPS-RSU +Ampakine** |  |
| 8h | min. | 13.75±1.96 | 180±0 | 41±4.59 |  |

**Table 2: Statistical analysis details**

|  | Test Used | | n | | Descriptive stats (Average, variance) | P Value | DEGREES OF  FREEDOM &  F/t/z/R/ETC VALUE |
| --- | --- | --- | --- | --- | --- | --- | --- |
| 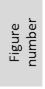 | Which test | 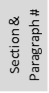 | Exact value | Defined | Reported | Exact Value | Value |
| 1 d | Two way RM ANOWA test, followed by a Tukey’s multiple comparison test | Fig. legend | 5, 6 | No. of Slices from  at least 3 rats/group | Error bars are mean +/- s.e.m. | P=0.011 | F(1,9)=10.16 |
| 1 e | Two way RM ANOWA test, followed by a Tukey’s multiple comparison test | Fig. legend | 5, 6 | No. of Slices from  at least 3 rats/group | Error bars are mean +/- s.e.m. | P<0.001 | F(1,9)=23.064 |
| 1 g | TW ANOWA test, followed by a Tukey’s multiple comparison test | Fig. legend;  Result para 1 | 6 in each group | No. of cells from  at least 3 rats/group | Error bars are mean +/- s.e.m. | P=0.009 | F(1,20)=8.437 |
| 1 i | TW ANOWA test, followed by a Tukey’s multiple comparison test | Fig. legend:  Result para 1 | 6 in each group | No. of Slices from  at least 3 rats/group | Error bars are mean +/- s.e.m. | P=0.0017 | F(1,20)=6.80 |
| 1 j | TW ANOWA test, followed by a Tukey’s multiple comparison test | Fig. legend;  Result para 1 | 6 in each group | No. of Slices from  at least 3 rats/group | Error bars are mean +/- s.e.m. | p-0.465 | F(1,20)=0.465 |
| 1 k | TW ANOWA test, followed by a Tukey’s multiple comparison test | Fig. legend:  Result para 1 | 6 in each group | No. of Slices from  at least 3 rats/group | Error bars are mean +/- s.e.m. | P=0.573 | F(1,20)=0.328 |
| 1 l | TW ANOWA test, followed by a Tukey’s multiple comparison test | Fig. legend;  Result para 1 | 6 in each group | No. of Slices from  at least 3 rats/group | Error bars are mean +/- s.e.m. | p-0.883 | F(1,20)=0.022 |
| 2 e | One way ANOWA by ranks, followed by a Tukey’s multiple comparison test | Fig. legend;  Result para 2 | 7 in each group | No. of cells from at least 3 different well plates per group | Error bars are mean +/- s.e.m | P=0.001 | H=13.588 with 2 degrees of freedom |
| 3 c | Student’s T- Test | Fig. legend;  Result para 3 | 9,12 | No. of rats per group | Error bars are mean +/- s.e.m | P<0.001 | T=17.357 |
| 5 c | TW ANOWA test, followed by a Tukey’s multiple comparison test | Fig. legend;  Result para 4 | 6 in each group | No. of Slices from  at least 3 rats/group | Error bars are mean +/- s.e.m | P<0.001 | F(1,20)=32.072 |
| 5 f | TW ANOWA test, followed by a Tukey’s multiple comparison test | Fig. legend;  Result para 5 | 8 in each group | No. of rats per group | Error bars are mean +/- s.e.m | P<0.001 | F(1,28)=127.762 |
| 5 g | TW ANOWA test, followed by a Tukey’s multiple comparison test | Fig. legend;  Result para 5 | 8 in each group | No. of rats per group | Error bars are mean +/- s.e.m | P<0.001 | F(1,28)=235.256 |
| 6 b | TW ANOWA test, followed by a Tukey’s multiple comparison test | Fig. legend;  Result para 6 | 8 in each group | No. of rats per group | Error bars are mean +/- s.e.m | P<0.001 | F(1,28)=1341.31 |
| 6 d | TW ANOWA test, followed by a Tukey’s multiple comparison test | Fig. legend;  Result para 6 | 8 in each group | No. of rats per group | Error bars are mean +/- s.e.m | P<0.001 | F(1,28)=3975.11 |
| 6 h | TW ANOWA test, followed by a Tukey’s multiple comparison test | Fig. legend;  Result para 7 | 7 in each group | No. of rats per group | Error bars are mean +/- s.e.m | P<0.001 | F(1,24)=47.046 |
| 7 c | One way ANOWA by ranks, followed by a Tukey’s multiple comparison test | Fig. legend;  Result para 8 | 6 in each group | No. of hippocampal tissue per group | Error bars are mean +/- s.e.m | P=0.010 | H=11.37 with 3 degrees of freedom |
| 7 d | TW ANOWA test, followed by a Tukey’s multiple comparison test | Fig. legend;  Result para 8 | 6 in each group | No. of hippocampal tissue per group | Error bars are mean +/- s.e.m | P=0.014 | F(1,20)=7.310 |
| 7 e | TW ANOWA test, followed by a Tukey’s multiple comparison test | Fig. legend;  Result para 8 | 6 in each group | No. of hippocampal tissue per group | Error bars are mean +/- s.e.m | P=0.002 | F(1,20)=13.266 |
| 7 h | One way ANOWA by ranks, followed by a Student-Newman –Keuls  multiple comparison test | Fig. legend;  Result para 8 | 12 in each group | No. of hippocampal tissue per group | Error bars are mean +/- s.e.m | P = 0.002 | H = 14.798 with 3 degrees of freedom. |
| 7 i | One way ANOWA by ranks, followed by a Student-Newman –Keuls  multiple comparison test | Fig. legend;  Result para 8 | 12 in each group | No. of hippocampal tissue per group | Error bars are mean +/- s.e.m | P <0.001 | H = 30.874 with 3 degrees of freedom. |
| 7 j | TW ANOWA test, followed by a Tukey’s multiple comparison test | Fig. legend;  Result para 8 | 12,10,11,10 | No. of slices from 3 to 5 rats/group | Error bars are mean +/- s.e.m | P<0.001 | F(1,35)=56.65 |
| 8 c | Mann Whitney test | Fig. legend;  Result para 9 | 5 | No. of cells from 3 rats/group | Error bars are mean +/- s.e.m | P=0.008 | T=15.00 |
| 8 e | One way ANOWA by ranks, followed by a Tukey’s multiple comparison test | Fig. legend;  Result para 9 | 5 in each group | No. of cells from 3 rats/group | Error bars are mean +/- s.e.m | P=0.004 | H=11.06 |
| 8 h | One way ANOWA test, followed by a Tukey’s multiple comparison test | Fig. legend;  Result para 9 | 7 in each group | No. of rats  per group | Error bars are mean +/- s.e.m | P<0.001 | F(2,18)=1106.708 |

**Table 3: Antibodies**

| **Antibody** | **Type** | **Dilution** | **Company** | **Catalog Number** | **Figure #** |
| --- | --- | --- | --- | --- | --- |
| **TLR4** | Rabbit Polyclonal | 1:500 | Santa Cruz | H-80 | Fig 1a |
| **TLR4** | Mouse Monoclonal | 1:1000 | Cell Signaling | 2219 | Fig 7b,c |
| **MAP2** | Chicken Polyclonal | 1:1000 | AbCam | ab5392 | Fig. 2a |
| **NeuN** | Mouse monoclonal | 1:1000 | Millipore | MAB377 | Fig. 1a |
| **GluA1** | Rabbit Polyclonal | 1:1000 | Millipore | MAB1504 | Fig 1h |
| **GluA2** | Mouse monoclonal | 1:1000 | Millipore | MAB397 | Fig 1h |
| **GFAP** | Mouse monoclonal | 1:1000 | Millipore | MAB360 | Fig 2b, 7b |
| **IBA-1** | Mouse monoclonal | 1:1000 | Millipore | MABN92 | Fig 2b, 7b |
| **β-actin** | Mouse monoclonal | 1:5000 | Sigma-Aldrich | A5441 | Fig 1h,7b |
| **CD45** | Mouse IgG1 | 1:50 | BD Biosciences | 740258 | Fig7f |
| **CD45 Isotype control** | Mouse IgG1 | 1:50 | BD Biosciences | 563547 | - |
| **CD4** | Mouse IgG2a | 1:50 | BD Biosciences | 550296 | Fig 7f |
| **CD4 Isotype** | Mouse IgG2a | 1:50 | BD Biosciences | 550339 | - |
| **CD3** | Mouse IgM | 1:50 | BD Biosciences | 557030 | Fig 7f |
| **CD3 Isotype Control** | Mouse IgM | 1:50 | BD Biosciences | 550883 | - |
| **Ox42** | Mouse IgG2a | 1:100 | BioLegend | 201820 | Fig 7f |
| **Ox42 Isotype Control** | Mouse IgG2a | 1:100 | BioLegend | 400258 | - |
| **Gr1** | Mouse IgG2a | 1:50 | BD Biosciences | 550002 | Fig 7f |
| **Gr-1 Isotype Control** | Mouse IgG2a | 1:50 | BD Biosciences | 553457 | - |
